# Structure and symmetries of the genetic codes for vertebrate, ascidian and yeast mitochondria

**DOI:** 10.1101/2023.07.23.548166

**Authors:** Rajeev Kohli

## Abstract

Genetic codes assign sixty-four codons to twenty amino acids. These assignments are known to follow certain rules. One question long considered but still unresolved is if these rules are derived from an underlying structure in genetic codes. Knowledge of such structure could facilitate better understanding of the biochemical, physico-chemical and evolutionary causes of the observed codon assignments. Our first finding reveals a coherent and symmetric structure in the genetic codes of vertebrate, ascidian, and yeast mitochondria. This structure is derived from a “simple” code that assigns all four codons with the same second nucleotide to a single amino acid if the first nucleotide is C or G, and assigns pairs of codons with the same second nucleotide to a single amino acid if the first nucleotide is A or U, and their third nucleotides are both purines (A and G) or both pyrimidines (U and C). The translation mechanism for the three mitochondria reflects this structure, one tRNA decoding each group of two or four codons into an amino acid. Our second finding is that the mycoplasma/spiroplasma and standard genetic codes are obtained by small sequential modifications of the vertebrate mitochondrial code and retain almost all its symmetries. We use group theory to characterize the symmetries of the simple and mitochondrial codes, and speculate on the implications of the structure for detecting translation errors and the evolution of the genetic code.

## Introduction

As the genetic code was deciphered, researchers such as Brenner, Crick, Khorana, Matthaei, and Nirenberg noticed patterns in the assignment of codons to amino acids. Crick^1^ described these patterns for the standard genetic code, which with some modifications are observed in other genetic codes that have been discovered since.^2–7^ Understanding the origins of these patterns has been a significant area of research for over sixty years.^8–18^

One question long considered but still unresolved concerns the existence of a structure that provides a simple and coherent account of the various observed patterns.^1, 25^ Many discussions conflate this question with another, which asks for the reasons for the observed codon assignments. But structure cannot replace the explanations that have been proposed for codon assignments in terms of their biochemical and physico-chemical properties, and the evolution and co-evolution of genetic codes.^12, 19^ Instead, it can help by focusing the explanations on the essential features of the structure.

Building on this foundation, our study uncovers a notable common structure in the genetic codes of vertebrate, ascidian, and yeast mitochondria (Figure 2). These three genetic codes display remarkable symmetry. Their structure can be derived from another “simple’ code” (Figure 1). In this simple code, all four codons with the same second nucleotide are assigned to the same amino acid if their first nucleotide is either C or G (family boxes); and pairs of codons with the same second nucleotide are assigned to the same amino acid if their first nucleotide is either A or U, and their third nucleotides are both purines (A and G) or pyrimidines (U and C) (two-codon sets). The structure of the three mitochondrial genetic codes (Figure 2) emerges by “reflecting” a subset of codon assignments in the simple code. This structure is related to the translation mechanisms of the three mitochondria, a single tRNA decoding a group of two, or four, codons into an amino acid.^2, 20–22^ Vertebrate mitochondria uses only 22 tRNAs,^3^ and yeast and ascidian mitochondria 24 each.^22, 23^

**Figure 1:**
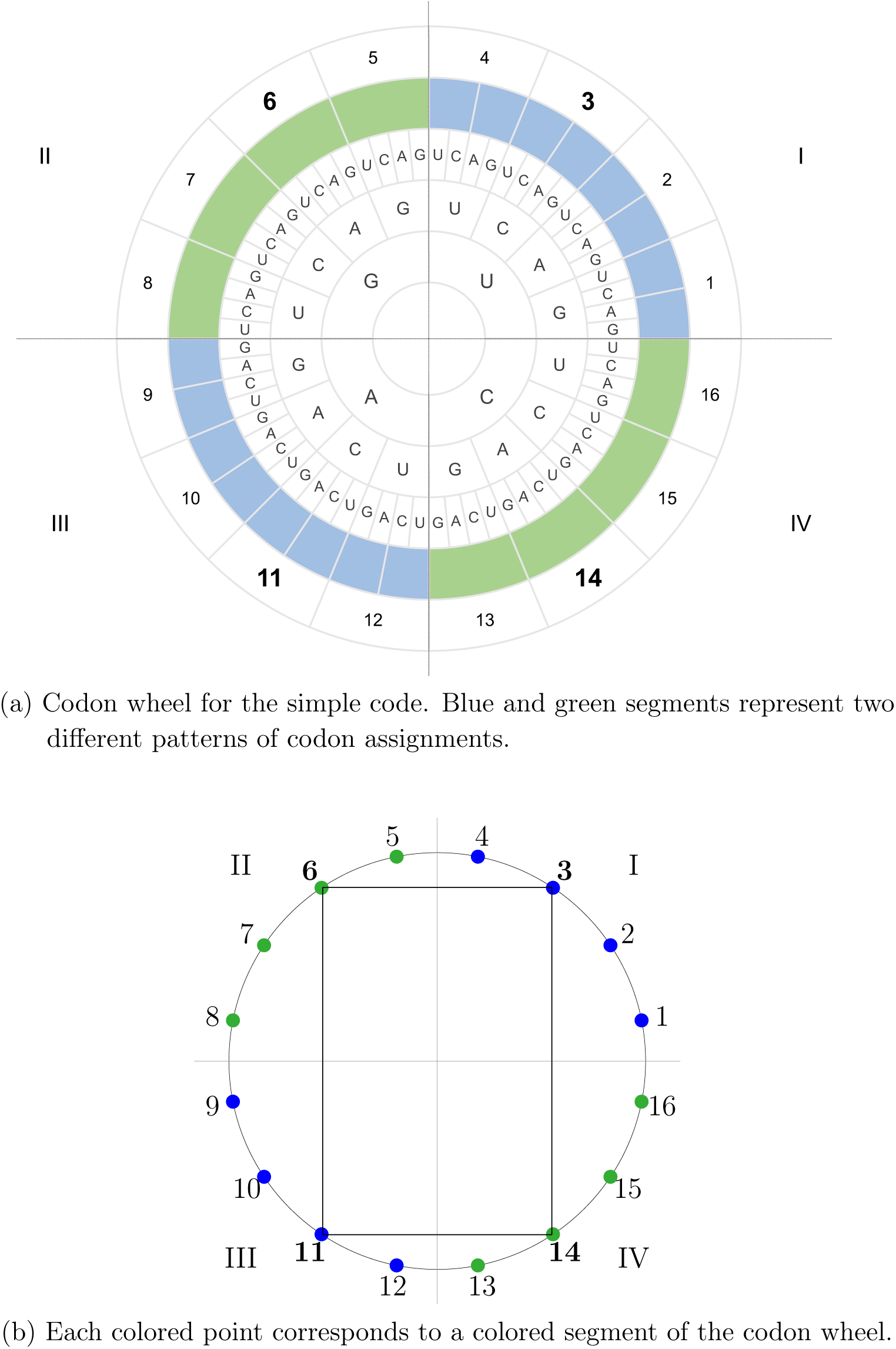
Simple code.

**Figure 2:**
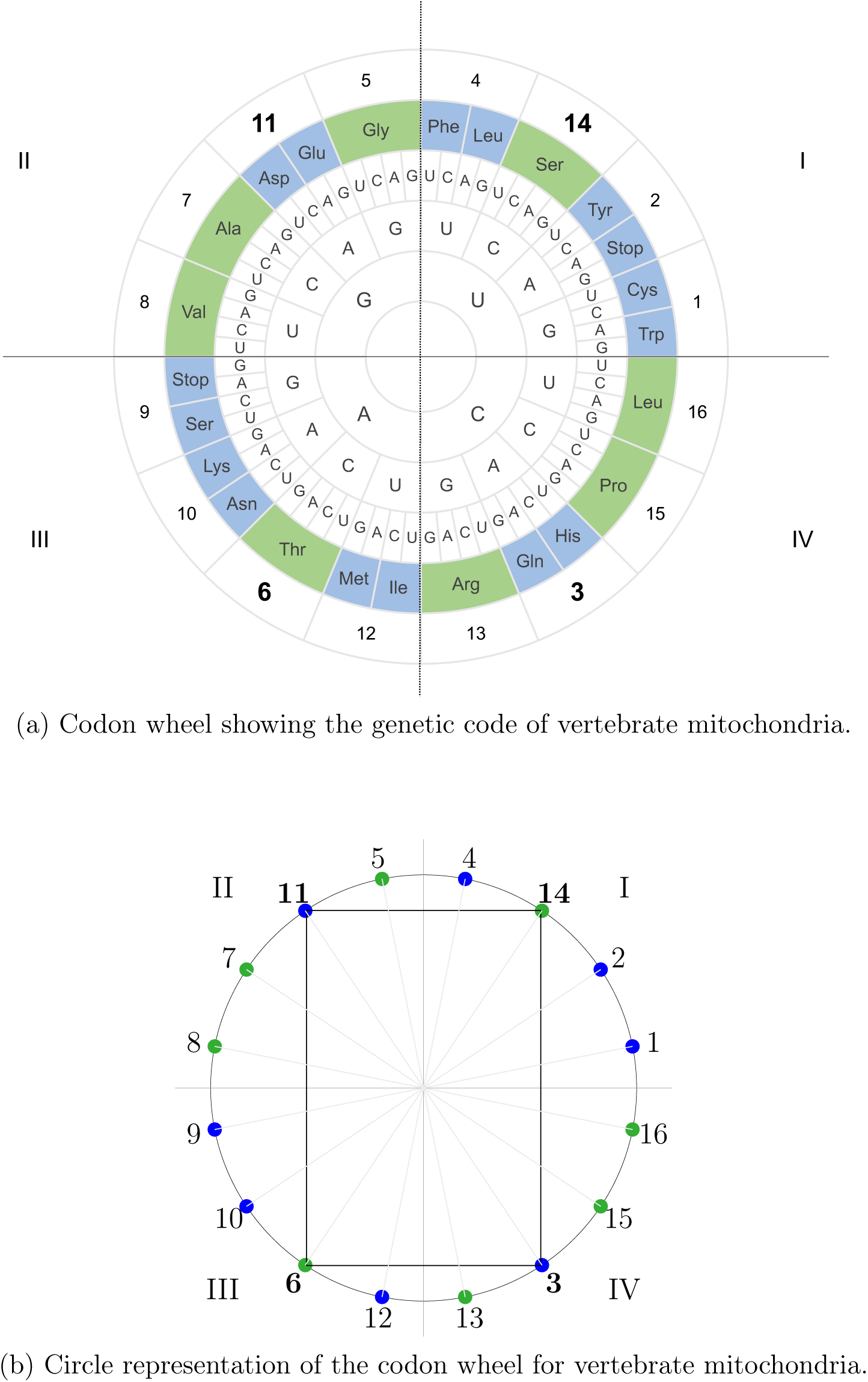
Genetic code of vertebrate mitochondria.

We use group theory to examine the symmetries of the simple and mitochondrial genetic codes. These symmetries are described by the transformations that preserve the structures of their codon to amino acid assignments. The symmetries are largely retained in the genetic code used by mold, protozoan and coelenterate mitochondria, and by mycoplasma and spiroplasma bacteria. They are also largely retained by the standard genetic code. Mycoplasma and spiroplasma also use a small number of tRNAs.^28–32^ For example, the *Mycoplasma capricolum* genome contains 30 tRNA genes for 29 tRNA species.^7^ However, organisms using the standard genetic code employ a much larger number of tRNAs to decode codons into amino acids. To our knowledge, these structures and relations have not been previously described, although group theory has been used to examine the structure and evolution of genetic codes.^9, 12, 14, 16, 24, 26, 27^

We speculate about the implications of the reported structures for error detection in the translation machinery, and for the evolution of these genetic codes. Regarding error detection, a violation of a symmetry can indicate an error in the translation machinery. Regarding code evolution, endosymbiotic theories propose that mitochondria evolved from bacteria.^4, 15^ Consistent with these theories, the aforementioned symmetries in the genetic codes and translation mechanisms could have originated in bacteria and transmitted in unaltered form to the three mitochondria. There is also substantial evidence that mycoplasma and spiroplasma bacteria evolved from larger bacteria.^26, 34^ Since the codon assignment structure for mycoplasma/spiroplasma differs only in one codon assignment from that of the three mitochondria, it is possible that the same common bacterial ancestor evolved along a different line into mycoplasma and spiroplasma. The standard genetic code could also have originated in bacteria with the same symmetric structure since its genetic code differs from that of mycoplasma/spiroplasma in a single codon assignment.

## Results

We describe a “simple code,” from which we derive the structure of codon assignments for the vertebrate, ascidian and yeast mitochondrial genetic codes. We relate this structure to tRNA assignments and use group theory to characterize its symmetries. The genetic code for vertebrate mitochondria differs only slightly from the mycoplasma/spiroplasma, protozoa, mold, and coelenterate mitochondrial genetic codes, which in turn differ only slightly from the standard genetic code. These codes retain almost all the symmetries of the three mitochondrial genetic codes.

### Simple code

Figure 1(a) shows the codon wheel for a “simple code.” It does not correspond to any known genetic code but has a notable and visible structure. We use the simple code to derive the (common) structure of the vertebrate, ascidian and yeast mitochondrial genetic codes.

Figure 1(a) divides the sixty-four codons into sixteen segments, each with four codons that share the same first and second nucleotides. The segments are numbered 1 to 16 in the anti-clockwise direction. Four segments appear in each quadrant I–IV. A segment is colored green if all four of its codons are assigned to one amino acid (a family box), and blue if pairs of codons terminating in purines (A, G) and pyrimidines (U, C) are assigned to different amino acids (two-codon sets). All segments in the same quadrant have the same color, and segments in adjacent quadrants have different colors. Equivalently, if XYZ denotes a codon, where X, Y, Z *∈* {U, C, A, G}, then:

1. All four codons XYU, XYC, XYA and XYG code for the same amino acid, where X*∈* {C, G} and Y *∈*{U, C, A, G}.
2. XYA and XYG code for one amino acid, and XYU and XYC code for another amino acid, where X*∈* {U, A} and Y*∈*{U, C, A, G}.

Figure 1(b) represents each segment of the codon wheel in Figure 1(a) by a numbered and colored point on a circle. We observe the following two features of Figure 1(b): (1) two points connected by a diameter of the circle have the same color; and (2) two points connected by a vertical or horizontal line have different colors. The first feature implies that the coloring pattern in Figure 1(b) is unchanged if we rotate all the points on the circle by 180 degrees. The second feature implies that if we “fold” the circle inward across the horizontal or vertical axis, each blue point aligns with a green point. That is, the coloring pattern is unaltered if we reflect all the points across the horizontal or vertical axis through the center of the circle and change their colors. The coloring pattern is also unchanged if we rotate the points on the circle by 90 degrees in the clockwise or anti-clockwise direction and change their colors.

The coloring pattern in Figure 1(b) implies that if we know the color of any single point on the circle, we can generate the colors of all other points by (1) assigning the same color to all other points in the same quadrant and in the diagonally opposite quadrant, and (2) assigning the other color to all points in the adjacent quadrants. For example, suppose we know that point 1 is colored blue (that is, UGG and UGA are assigned to one amino acid, and UGC and UCU to another amino acid). Then we can assign blue to all other points in quadrants I and III, and green to all points in quadrants II and IV.

The code in Figure 1 can be itself derived from a still simpler code in which all four codons that share the same first two codons are assigned to a single amino acid. Jukes^35^ and Osawa et al.^7^ suggested that such a code might have been present in an earlier (archetypal) form of the genetic code. The simple code could be obtained from it by a symmetric division of the family boxes starting with the A and U into two-codon sets. Below, we consider how the simple code is related to the genetic codes of vertebrate, ascidian and yeast mitochondria.

### Codes for vertebrate, ascidian and yeast mitochondria

Observe that the points 3, 6, 11 and 14 form the vertices of a rectangle in Figure 1(b) and have alternating blue and green colors. The codon assignment pattern for the vertebrate, ascidian and yeast mitochondria is obtained by reflecting (exchange) the pairs of points 3 and 14, and 6 and 11 across the horizontal axis. Figure 2(b) shows the resulting colored wheel. Figure 2(a) shows the corresponding codon wheel, as well as the genetic code for vertebrate mitochondria. If XYZ denotes a codon, then Figure 2 displays the following pattern of codon assignments:

1. Let X*∈* {C, G}. Then all four codons XYU, XYC, XYA and XYG code for the same amino acid when Y *∈*{U, C, G}; and XYA and XYG code for one amino acid, and XYU and XYC code for a different amino acid, when Y=A.
2. Let X*∈* {U, A}. Then XYA and XYG code for one amino acid, and XYU and XYC code for a different amino acid, when Y*∈*{U, A, G}; and all four codons XYU, XYC, XYA and XYG code for the same amino acid when Y=C.

The coloring in Figure 2 is also obtained by reflecting the points 3, 6, 11 and 14 across the vertical axis, instead of the horizontal axis, in Figure 1. And the colorings in Figures 1 and 2 are complementary: reflecting the points 3, 6, 11 and 14 across the horizontal or vertical axis in one figure produces the coloring in the other.

All the points in Figure 2(b) can be colored if we know the position and color of the “odd” point in any one quadrant. For example, suppose we know that the green point numbered 14 is the “odd” point in quadrant I. Then we can color all other points in quadrant I blue. Next, we draw horizontal and vertical lines from each point in quadrant I and identify the points where these lines intersect the circle in quadrants II and IV. If a point in quadrant I is colored blue, we color the intersecting points in quadrants II and IV green; and if it is colored green, we color the intersecting points in quadrants II and IV blue. Finally, we draw diameters through each point in quadrant I and assign the same color to each diametrically opposite point in quadrant III.

Table 1 shows the differences in the codon assignments for the genetic codes of vertebrate, yeast and ascidian mitochondria. The genetic code for vertebrate mitochondria uses the codons AGA and AGG as termination signals, whereas the genetic code of ascidian mitochondria uses these two codons to represent glycine. And the genetic code for yeast mitochondria differs from the genetic code for ascidian mitochondria by assigning the AGA and AGA codons to arginine instead of glycine, and the CUU, CUC, CUA and CUG codons to threonine instead of leucine. However, none of these differences change the pattern of codon assignments (the coloring) in Figure 2.

**Table 1:** Differences in the amino acids assigned to codons in the vertebrate, ascidian and yeast mitochondrial genetic codes.

| Codons | Vertebrate | Ascidian | Yeast |
| --- | --- | --- | --- |
| CUU, CUC, CUA, CUG | Leucine | Leucine | Threonine |
| AGA and AGG | Stop | Glycine | Arginine |

Note: Amino acid assignments that change between successive genetic code are in red.

### Symmetries of the genetic codes

The symmetries of the simple code and the three mitochondrial genetic codes are described by transformations that leave the colorings of the points in Figures 1(b) and 2(b) unchanged. At the broadest level, these transformations allow (1) exchanging the positions of any two points that have the same color, and (2) exchanging any two points that have different colors and reversing the colors of the two points. These transformations include the following two, which were previously noted for the simple code and are unaltered in the genetic codes for the three mitochondria. Referring to Figure 2(b), two points connected by a diameter of the circle have the same color (Figure 3), and two points connected by a vertical or horizontal line have different colors. The latter implies that the four rectangles shown in Figure 4 have vertices with alternating colors. Thus, the coloring pattern in Figure 2(b) is unchanged by (1) a 180-degree rotation, and (2) a reflection of the points across the horizontal (or vertical) axis combined with change in the color of all points on the circle. This group of symmetries is isomorphic to the Klein four-group^41^ (see the Methods section).

**Figure 3:**
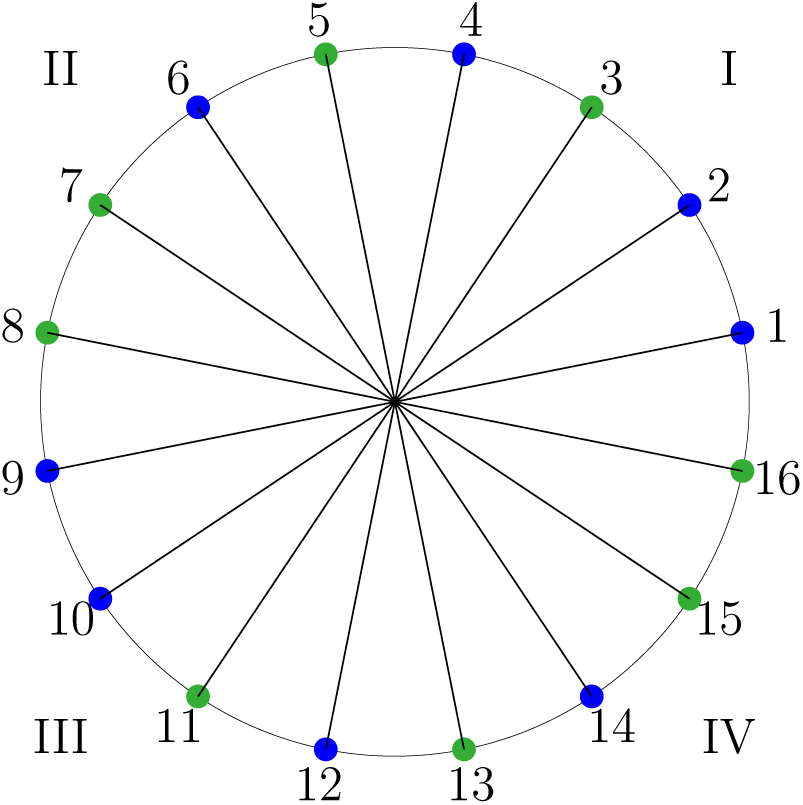
Diameters of the circle representing the genetic code for the three mitochondria connect points with the same color.

**Figure 4:**
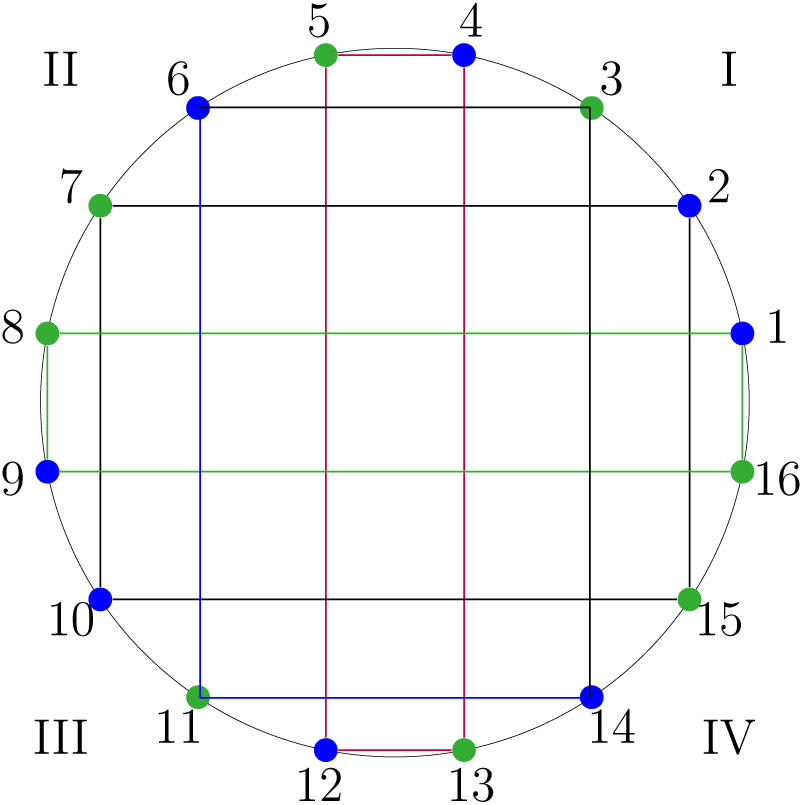
Rectangles inscribed in a circle representing the genetic codes for the three mitochondria have vertices with alternating colors.

Next, we consider symmetries of the simple code that are changed when it is modified to produce the pattern of codon assignments in the genetic codes for the three mitochondria. Unlike in Figure 1(b), the coloring of the circle in Figure 2(b) is changed by a 90-degree clockwise or anti-clockwise rotation combined with a change in color for all points on the circle. The reason is that Figure 2(b) has one odd-colored point in each quadrant. However, the colorings in both Figures 1(b) and 2(b) are unaffected by a 90-degree rotation combined with both an involution and a color change of all the points on each circle. (An involution means inverting the ordering of points in a quadrant.) For example, consider the points 1, 2, 14 and 4 in quadrant I of Figure 2(b). A 90-degree rotation in the anti-clockwise direction brings these points to quadrant II; an involution reverses the ordering of these points to 4, 14, 2, 1; and reversing the coloring of these points yields the green-blue-green-green color sequence associated with the points 5, 11, 7 and 8 in Figure 2(b).

Second, in Figure 1(b), any two points connected by lines (including horizontal and vertical lines) in adjacent quadrants have opposite colors. Consequently, exchanging two such points and changing their colors does not alter the coloring of the circle. This property still holds for points that have different colors in adjacent quadrant in Figure 2(b). But since each quadrant has one odd-colored point, there are only ten such pairs of points in adjacent quadrants. The other six pairs of points in adjacent quadrants have the same color and can be reflected without changing their colors.

For example, Figure 5 shows the four squares that can be inscribed in the circle in Figure 2(b). One square connects only blue vertices, another only green vertices, and the other two alternating blue and green vertices. In the first two cases, the coloring of the circle is unchanged by reflecting any two vertices in adjacent quadrants without changing their colors; in the latter two cases, it is unchanged if reflecting vertices in adjacent quadrants is accompanied by changing their colors.

**Figure 5:**
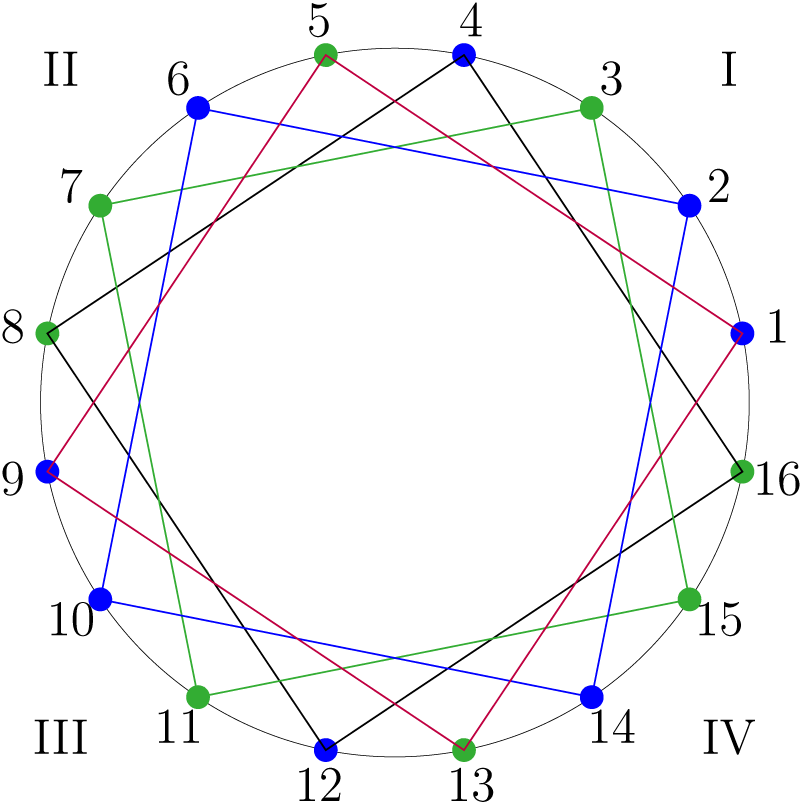
Squares 2-6-10-14 and 3-7-11-15 have all blue and all green vertices. Squares 1-5-9-13 and 4-8-12-16 have vertices with alternating colors.

### The mycoplasma/spiroplasma and standard genetic codes

The preceding pattern of assignments for the three mitochondrial genetic codes is slightly modified in the genetic code common to mycoplasma/spiroplasma bacteria, protozoa, mold, and coelenterate mitochondria. And the latter code is slightly modified in the standard genetic code. Table 2 shows the differences between these and the vertebrate mitochondrial genetic code. The first difference, which does not change the coloring (codon assignment pattern) in Figure 2(b), is that the mycoplasma/spiroplasma genetic code assigns the AGA and AGG codons to arginine instead of using them as termination signals. The second difference, which does alter the coloring pattern in Figure 2(b), is that the mycoplasma/spiroplasma genetic code assigns the AUA codon to isoleucine instead of methionine. Table 2 also shows that the standard and mycoplasma/spiroplasma genetic codes differs only in the reassignment of the UGG codon to a terminating signal instead of tryptophan. Thus, two small, successive, changes link the vertebrate mitochondrial genetic code first to the mycoplasma/spiroplasma genetic code and then to the standard genetic code.

**Table 2:** Successive differences in the amino acids assigned to codons in the vertebrate, mold and standard genetic codes.

| Codons | Vertebrate mitochondrial | Mycoplasma and Spiroplasma* | Standard |
| --- | --- | --- | --- |
| AGA, AGG | Stop, Stop | Arginine, Arginine | Arginine, Arginine |
| AUA, AUG | Methionine, Methionine | Isoleucine, Methionine | Isoleucine, Methionine |
| UGA, UGG | Tryptophan, Tryptophan | Tryptophan, Tryptophan | Stop, Tryptophan |

\* Also, protozoa, mold, and coelenterate mitochondria.
Note: Amino acid assignments that change between successive genetic code are in red.

### tRNA symmetries

The translation mechanisms for the three mitochondria display superwobbling^39, 40^ and closely reflect the common structure of their genetic codes. Vertebrate mitochondria use only 22 tRNAs.^2^ For each green segment coding for an amino acid in Figure 2(a), an unmodified uridine in the first (wobble) position of the anticodon recognizes all four nucleotides in a codon’s third position; and for each blue segment coding for two amino acids (excluding the two pairs of codon signaling termination), an unmodified uridine recognizes only pyrimidines, and a modified uridine discriminates purines from pyrimidines, in a codon’s third position.

The reassignment of AGA and AGG to glycine in the genetic code for ascidian mitochondria,^23^ and to arginine in the genetic code for yeast mitochondria, slightly increases the number of tRNAs. For instance, the mitochondrial genome of the species Saccharomyces cerevisiae (baker’s yeast) encodes 24 tRNAs, just enough to recognize each amino acid.^22^ The mitochondrial genomes for the ascidian species Halocynthia roretzi (sea pineapple) also has 24 tRNAs.^23^

Table 2 shows that the only difference between the genetic codes for vertebrate mitochondria and mycoplasma/spiroplasma bacteria is that the latter assigns the AUG codon to isoleucine, and the AGA and AGG codons to arginine. The translation mechanism for mycoplasma also displays superwobbling but to a lesser extent than for vertebrate mitochondria. For example, *M. capricolum*, *U. urealyticum* and *M. genitalium*/*M. pneumoniae* have 28, 29 and 32 anticodons, respectively.^28–30, 32^ Similarly, *Spiroplasma taiwanense*, *Spiroplasma diminutum* and *Spiroplasma chrysopicola* have 28, 28 and 32 tRNAs decoding the standard 20 amino acids.^31^ In contrast, the human genome encodes more than 500 tRNA genes, although nearly half of them are silent.^36^

## Discussion

The genetic codes for vertebrate, ascidian and yeast mitochondria share a symmetric structure (Figure 2) that can be derived by reflecting four subsets of codons in a simple code (Figure 1). The translation mechanisms for these mitochondria reflect the symmetric structures of their genetic codes; and translation in mycoplasma and spiroplasma displays superwobbling to a slightly lesser extent. We used group theory to characterize the symmetries of the three mitochondrial genetic codes. Almost all this symmetry is retained by the genetic codes of mycoplasma and spiroplasma bacteria, protozoa, mold, and coelenterate mitochondria, as well as the standard genetic code.

These symmetries can facilitate verification of the accuracy of a translation mechanism. For example, consider the translation of codons to amino acids in vertebrate mitochondria. Recall that each green point is “decoded” using one tRNA and each blue point using two tRNAs. Since the four vertices of each rectangle in Figure 4 have alternating colors, the number of tRNAs used to decode a group of four codons around a rectangle should form an alternating sequence of ones (green points) and twos (blue points). A violation of this condition in any rectangle implies a translation error. However, while feasible, we do not know if such checks are used by the translation machinery.

Endosymbiotic theories propose that mitochondria evolved from bacteria.^4, 15^ Consistent with these theories, the highly symmetric structure of the genetic codes (and translation mechanisms) for vertebrate, ascidian and yeast mitochondria could be descended from bacteria. Support for this possibility also comes from the close relation between these three mitochondrial codes and the genetic code of mycoplasma and spiroplasma bacteria^33^ (Table 2). There is substantial evidence that mycoplasma and spiroplasma evolved from the ancestors of gram-positive bacteria by several rounds of genome reduction.^26, 34^ The genetic code of these ancestor bacteria could have had the same symmetries that were passed unchanged to vertebrate, ascidian and yeast mitochondria in one direction, and passed with slight modification to mycoplasma and spiroplasma bacteria, and to protozoa, mold, and coelenterate mitochondria, in another direction. The high degree of similarity between the mycoplasma/spiroplasma genetic code and the standard genetic code (Table 2) suggests that the latter might also have a bacterial ancestor with the symmetries shown in Figure 2. This could have occurred without a direct evolutionary path between the mycoplasma/spiroplasma genetic code and the standard genetic code.

Osawa et al.^7^ observed that family boxes are the simplest, perhaps the earlier, “units” in the genetic code. Jukes^35^ suggested that family boxes alone might have been present in an earlier (archetypal) form of the genetic code. In this case, the simple code could have been obtained by a symmetric division of half the family boxes (those starting with the A and U) into two-codon sets. And then, as we have discussed, the structure of the genetic codes for vertebrate, ascidian and yeast mitochondria (Figure 2) can be obtained by a reflection of a rectangle in the simple code (Figure 1).

## Methods

We used data on the genetic codes compiled by Elzanowski and Ostell^5^ at National Center for Biotechnology Information (NCBI) in Bethesda, Maryland, USA. The primary sources for these genetic codes are the reviews by Jukes and Osawa^6^ and Osawa et al.^7^ Elzanowski and Ostell^5^ used the DNA alphabet to describe genetic codes. We describe all genetic codes using the RNA alphabet, converting each T to U. We obtained information on spiroplasma tRNAs using the tRNAscan-SE search tool GtRNAdb.^31^ We used the cited sources to obtain information on all other tRNA assignments.

The Klein four-group^41^ describes the key symmetries shared by the simple code and the three mitochondrial genetic codes. It is the symmetry group of a non-square rectangle. The group has four elements, each element is its own inverse, and the composition of any two non-identity elements produces the third element. Let *G* = *{E, D_g_, D_b_, R}* denote the group of symmetric transformations for a rectangle in Figure 4. Here, (1) *E* denotes the identity transformation that does not change the coloring of a rectangle; (2) *D_g_* reflects the points across the diagonal from one green vertex to another, and *D_b_* across the diagonal from one blue vertex to another; and (3) *R* denotes the rotation of a rectangle by 180 degrees. It has the following multiplication table:

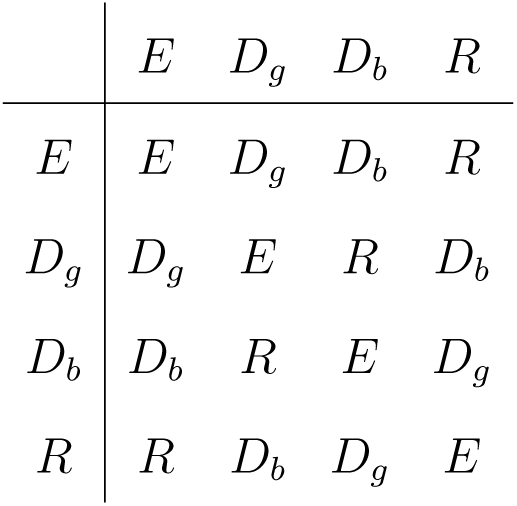

For example, the operation *D_g_* followed by *D_b_* gives *R* because reflecting a rectangle first across the *D_g_* diagonal (the one connecting green vertices), and then across the *D_b_*diagonal (the one connecting blue vertices) has the same effect as rotating the rectangle by 180 degrees.

## Notes

### Competing Interest Statement

The authors have declared no competing interest.

